# The superior colliculus arbitrates whole-brain dynamics for unconscious visual insight

**DOI:** 10.1101/2025.04.09.648050

**Authors:** Tsutomu Murata, Komaki Kunishige, Shigeto Seno, Izumi Ohzawa, Kunihiko Kaneko, Toshio Yanagida, Masahiko Haruno, Kazufumi Hosoda

## Abstract

The human capacity for sudden insight, often marked by abrupt and illuminating “eureka” moments, has long intrigued neuroscientists seeking to understand its elusive neural basis^1–6^. Mooney image recognition offers a compelling behavioral paradigm of this phenomenon, in which ambiguous black-and-white patterns are abruptly perceived as coherent objects^7–13^. Whole-brain cortical involvement, which is mainly revealed by the contrast between recognition and nonrecognition, is well established^8,14–27^. However, the neural dynamics underlying the transition from nonrecognition to recognition, particularly the contributions of noncortical structures, have remained largely unknown. Here, we show that the transition in whole-brain dynamics can be effectively characterized by three large-scale activity patterns and that the superior colliculus (SC) emerges as a structure with a distinct temporal profile that is potentially critical for insight. Using functional clustering of functional magnetic resonance imaging data comprising 41,446 time points from 14 human participants performing this insight task, we identified three distinct functional clusters exhibiting stimulus-driven activation, suppression, and recognition-associated patterns. Notably, the SC in the third cluster displayed a distinct activation peak immediately before the recognition response. Furthermore, empirical dynamic modeling revealed that the SC was the only region in the whole brain that exhibited mutual positive directed interactions with all three clusters and was functionally embedded within higher-order cortical interactions mediating the transformation from stimulus input to motor output. Our findings provide compelling experimental evidence that the SC plays a critical role in orchestrating whole-brain dynamics for sudden insight, calling for a reappraisal of this evolutionarily conserved structure as a key player supporting unconscious but high-level cognitive processing.

## Main text

Neuroimaging studies have demonstrated that resolving the ambiguity of Mooney images involves widespread cortical networks, including frontoparietal and default mode regions^8,14–24^. These neural processes have been investigated primarily using top-down guided paradigms involving pre-exposed undegraded images or forced-choice response formats, which are known to differ qualitatively from spontaneous recognition in open-ended tasks without prior information^4,13,28–32^.

In the absence of prior information, recognition has been associated with altered connectivity between visual and frontal regions^25–27^. Subcortical contributions to memory and reward processing have been reported^33,34^. The midbrain, thalamus, and cerebellum have also been implicated in other insight-related paradigms^35,36^. The SC, in particular, has been specifically discussed as a potential contributor in this context^19,37^. Traditionally linked to oculomotor control and unconscious visual processing^38–43^, the SC has more recently been implicated in higher-order functions such as attention and decision-making in forced-choice paradigms in nonhuman species^41,44–54^. However, its potential involvement in insight-related cognition and its integration within whole-brain dynamics remain unknown.

Here, we used functional magnetic resonance imaging (fMRI) to conduct a whole-brain neural time series analysis during an open-ended Mooney image recognition task performed without prior information. Although the visual stimulus remained unchanged throughout the task, widespread activation emerged around the moment of recognition. Notably, this data-driven analysis revealed a unique activity pattern in the SC, which stood out in comparison with all other brain regions.

### Mooney image recognition task without prior information

The participants viewed a series of black-and-white images and were instructed to press a button (response) upon recognizing the content of each stimulus (Figure 1A). To extract whole-brain dynamics, We covered 122 atlas-defined brain regions provided by SPM12^55,56^, excluding areas that did not contain neuronal tissue, to extract whole-brain dynamics. Within each atlas region, we performed prefunctional parcellation based on the voxel-wise correlations of time series concatenated across all participants (totaling 41,446 seconds) and performed standard first-level (task-based GLM for individual data) and second-level (group-level) analyses to exclude regions that were not related to the task. A total of 148 such subregions were retained for analysis, with each representative time series showing significant stimulus-related activity and collectively covering 64% of the neuronal tissue across the defined whole-brain atlas. These 148 representative time series formed the basis for subsequent whole-brain analyses (Figure 1B; see the Methods for full details).

**Figure 1.**
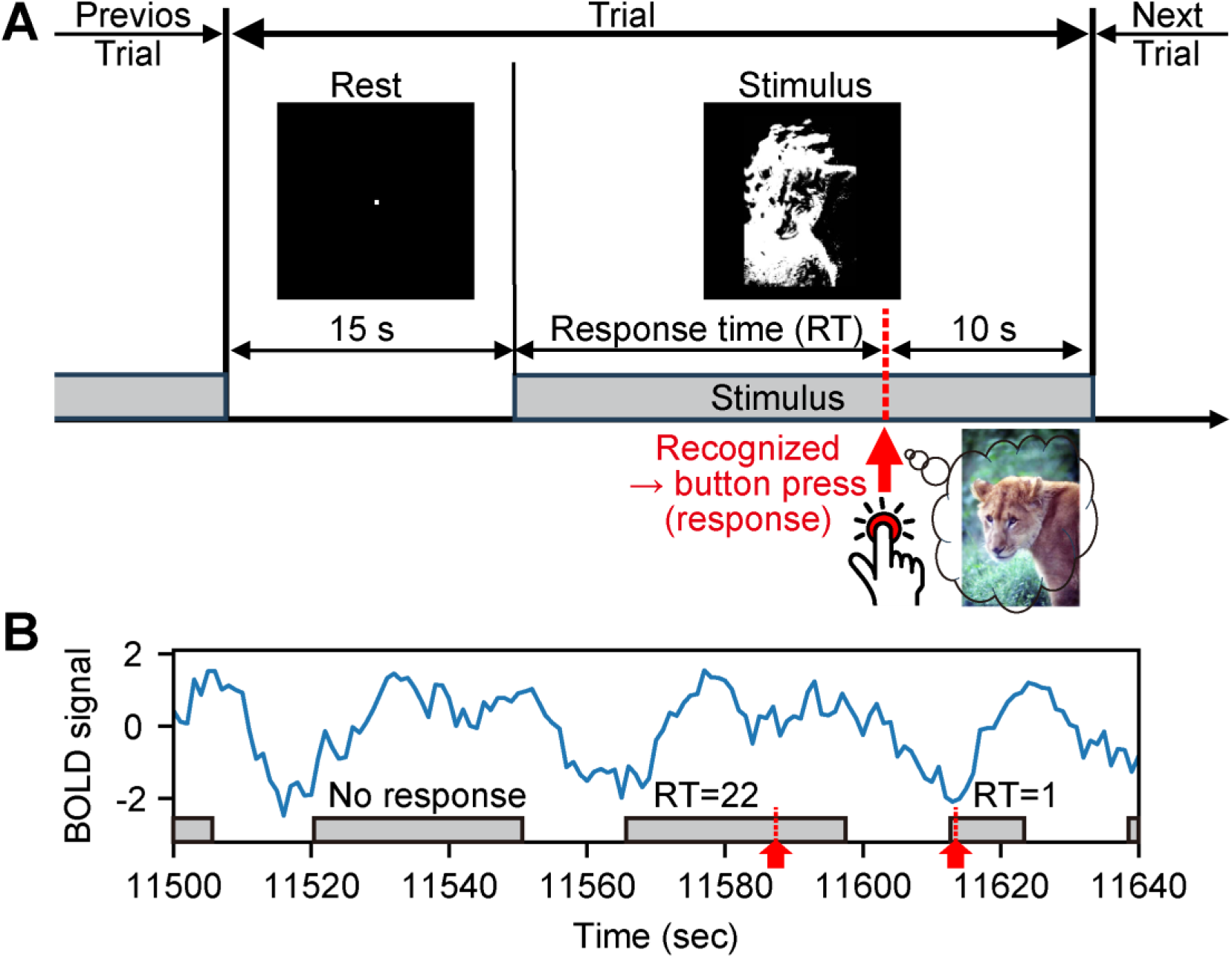
Experimental design and obtained time series data. **A**, Schematic of the experimental paradigm. The gray boxes indicate periods during which the stimuli were displayed, and the red arrow indicates the moment of the button press response. **B**, Example of the obtained time series data. The gray boxes and red arrows have the same meanings as those in Panel A. RT = 22 indicates that the response time, measured from stimulus onset, occurred between 22 and 23 seconds.

### Temporal organization of whole-brain activity

We examined trial-averaged time series from 148 functional subregions to obtain an overview of the whole-brain temporal structure during Mooney image recognition and found that many subregions exhibited similar temporal profiles (Figure 2A; trials with a response time (RT) greater than 18 seconds). Dimensionality reduction of the full 41,446-point time series using principal component analysis (PCA), followed by Gaussian mixture model (GMM) clustering, revealed that a three-cluster solution most efficiently captured the data structure (Figure 2B).

**Figure 2.**
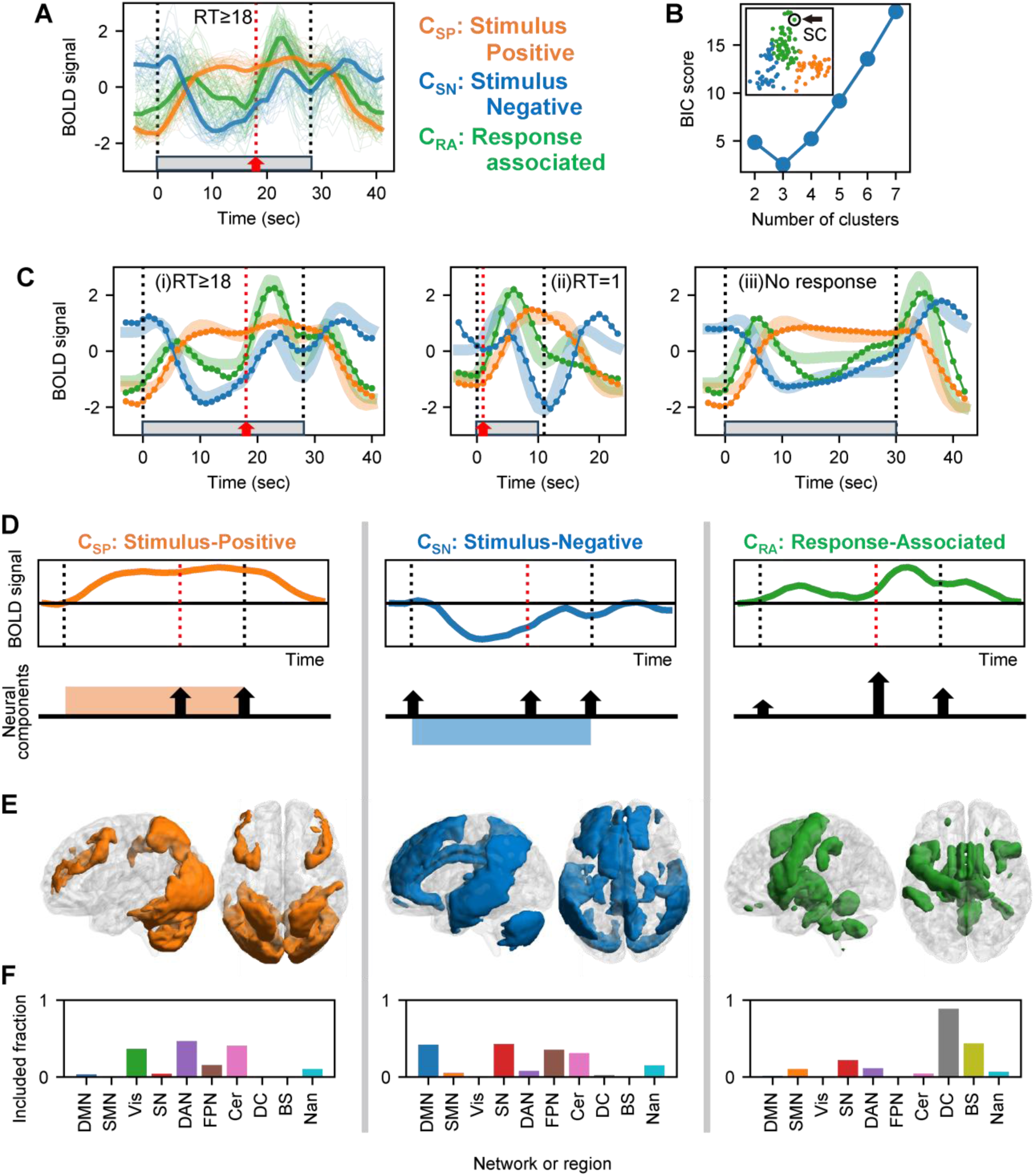
Overview of temporal dynamics. **A**, Time series of all 148 subregions across the brain. Thin lines represent individual subregions, color-coded by functional cluster identity (orange: C_SP_; blue: C_SN_; green: C_RA_). The thick lines represent the average time course within each functional cluster. The gray boxes and red arrows have the same meanings as those in Figure 1. The trials in which the response occurred after 18 seconds were aggregated under the assumption that these trials share similar BOLD time series both before and after the response (designated RT ≥ 18; N = 44). See the Methods for details. **B**, Bayesian information criterion (BIC) scores for Gaussian mixed models with varying numbers of clusters. The inset shows the clustering results for three clusters, with each of the 148 subregions projected into the PC1–PC2 state space. The superior colliculus (SC) can be seen as an edge within C_RA_. **C**, Model-based explanation of the time series dynamics. The model includes four event-related components, namely, stimulus onset, response, stimulus offset, and sustained stimulus presentation, each convolved with the canonical HRF. (i) Fit to the data from trials with RT ≥ 18. (ii, iii) Reconstructions of RT = 1 (N = 352) and no-response (N = 201) trials using the same fitted parameters without refitting. **D**, Summary of the temporal profile. The colored box shows the stimulus presentation, and the arrows show the contributions of each cluster to the three transient events. **E**, Spatial distribution of the three functional clusters. **F**, Proportion of each known large-scale brain network, as well as the diencephalon and brainstem regions included in the three functional clusters. For example, the value of 44% for the brainstem in the C_RA_ does not mean that 44% of the C_RA_ consists of brainstem regions but indicates that 44% of the brainstem voxels fall within the C_RA_, which corresponds to the entirety of the significantly prescreened brainstem mask. DMN, default mode network; SMN, sensorimotor network; Vis, visual network; SN, salience network; DAN, dorsal attention network; FPN, frontoparietal network; Cer, cerebellum; Thal, thalamus; DC, diencephalon; BS, brainstem. The CONN functional connectivity toolbox includes the language network in addition to the others, but since it is often considered part of the DMN, we treated it as part of the DMN in this study as well.

The resulting three functional clusters revealed interpretable patterns in the temporal dynamics of brain activity (Figure 2A). The first cluster, referred to as C_SP_ (stimulus-positive cluster), exhibited sustained high activation throughout the stimulus presentation, with a small transient peak around the recognition response. The second cluster, C_SN_ (stimulus-negative cluster), showed overall suppression during the stimulus period but was accompanied by an increase at the time of response. The third cluster, C_RA_ (response-associated cluster), displayed transient responses that peaked after key events, including stimulus onset, response, and stimulus offset. This temporal pattern aligned with a canonical hemodynamic delay and was most selectively engaged at the time of recognition. Importantly, all three clusters demonstrated a transient increase in activity at the moment of the recognition response, suggesting that this cognitive event elicits coordinated neural engagement across multiple functional subsystems.

We assessed how well the temporal profiles of the three functional clusters could be explained by discrete task events by constructing a model comprising four event-related components: stimulus onset, response, stimulus offset, and sustained stimulus presentation. Each component was convolved with the canonical hemodynamic response function (HRF), and the model was fitted separately to the average time course of each functional cluster (Figure 2C(i)). The fitted parameters were then used to simulate time courses for the different experimental conditions without any further fitting (Figure 2C(ii),(iii)), which captured the key features of the observed dynamics. A conceptual summary of these modeling results is shown in Figure 2D.

Anatomically, C_SP_ was distributed across the occipital and parietal lobes and the middle frontal gyrus, overlapping primarily with the visual (Vis) and dorsal attention (DAN) networks, as defined by the CONN functional connectivity toolbox^57^ (Figure 2E, F; visualized using BrainNet Viewer^58^). This pattern is consistent with sustained sensory engagement throughout stimulus presentation. The division between the cerebellar regions included in this cluster and those included in the following C_SN_ also aligned well with previously reported findings^59^. C_SN_ was found mainly in the temporal and frontal cortices, the cingulate gyrus, and the precuneus and corresponded closely to the default mode and salience networks. C_RA_ encompassed the diencephalon and brainstem, as well as cortical regions, including the anterior insula, frontal operculum, inferior frontal gyrus, supplementary motor cortex, and left primary motor cortex associated with the responding hand. This cluster showed peak activity at the moment of recognition, which suggested that recognition was supported not only by cortical networks but also by deep noncortical structures.

### The extraction of early response-related subregions highlights the superior colliculus

As the recognition response was accompanied by transient activation across nearly all subregions, we focused on identifying those that exhibited early activation. Specifically, we selected subregions whose 95% CI for the peak response occurred within 5 seconds after the response, based on trials with RTs greater than 8 seconds (Figure 3A).

**Figure 3.**
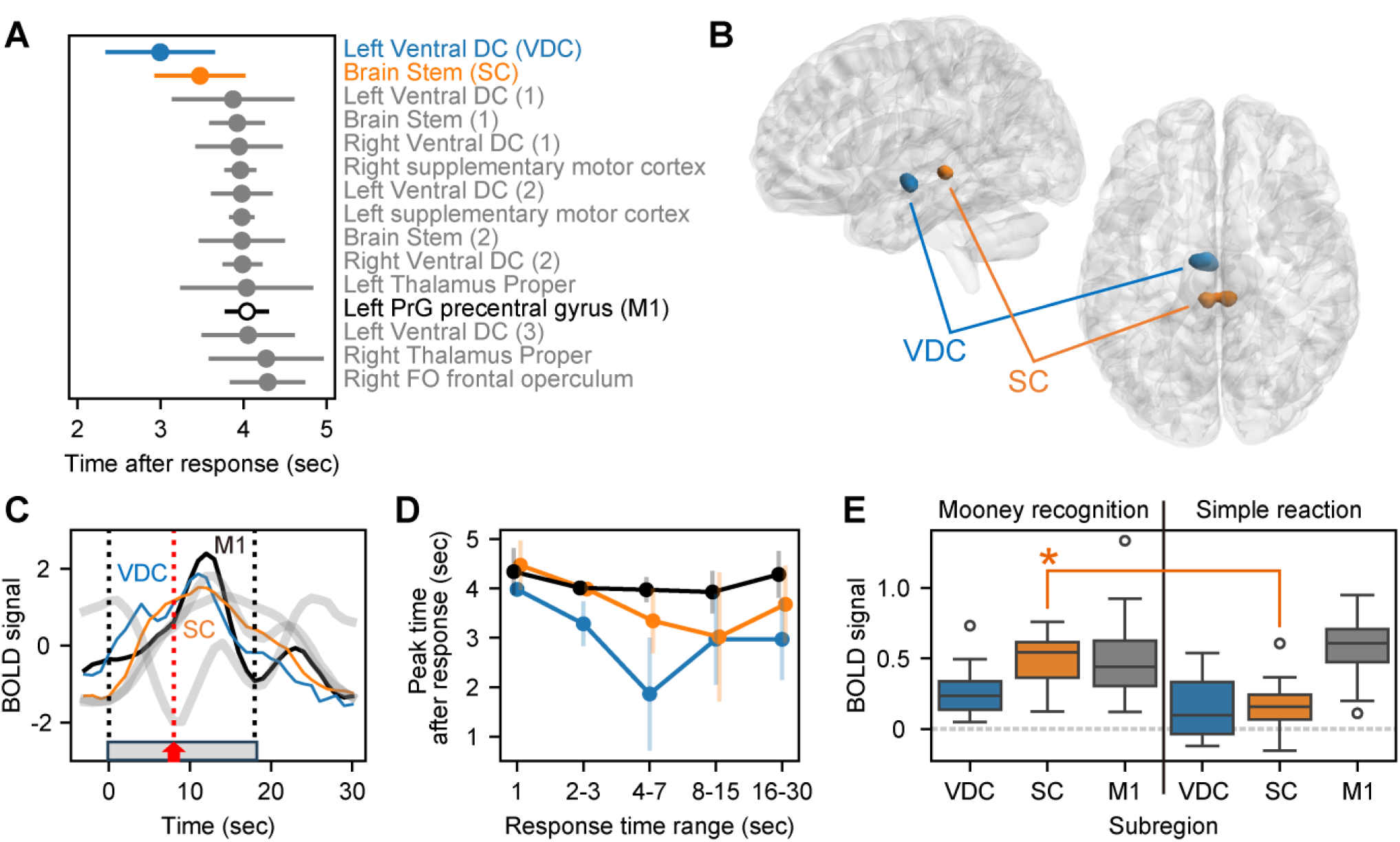
Subregions characterized by peak activation at the moment of the response. **A**, Subregions showing early BOLD signal peaks following the response. The displayed names correspond to the labels defined in the Neuromorphometrics atlas provided in SPM12. Multiple subregions may originate from the same atlas region, resulting in repeated label names, which are numbered accordingly; however, the voxel territories of the individual subregions do not overlap. The color coding for the top two subregions and the left precentral gyrus (M1) is consistent across all panels below. **B**, Anatomical locations of the top two subregions identified in Panel A. The brainstem region was identified as the SC. **C**, Time courses of the BOLD signals for the top two subregions and the left M1, averaged across trials with RT ≥ 8 seconds. The gray bold lines indicate the three functional clusters used for comparison. All time courses were standardized for display. **D**, Peak timing of the three subregions across different RT conditions. **E**, Comparison of BOLD signal amplitudes between the Mooney image recognition task and the simple reaction task for the three subregions. The data are shown as box plots across 14 participants. Orange asterisks indicate significant differences between the tasks. VDC, ventral diencephalon (top 1); SC, superior colliculus (top 2 brainstem); M1, left precentral gyrus. The error bars in all panels indicate the standard deviation.

All of these subregions belong to C_RA_. These subregions included the left primary motor cortex (M1), as expected for a right-hand motor response, as well as the bilateral supplementary motor area, known to be involved in preparation for self-initiated movement^60^. Most regions are located within the brainstem or diencephalon, which includes the thalamus; recently, the contribution of the thalamus to conscious perception has been revealed^61^.

Notably, two subregions displayed even earlier peak activity than M1 did (the 95% CI did not overlap with M1), one of which was anatomically localized to the superior colliculus (SC) based on a detailed brainstem atlas^62^ (Figure 3B). Notably, despite the absence of any prior use of atlas-based constraints for the SC, the region was highlighted with remarkable clarity and specificity (this 60-voxel subregion encompasses 90% of the SC, as defined by the brainstem atlas).

The temporal dynamics of these two subregions are shown in Figure 3C. M1 peaked earlier than the average of its associated cluster, and indeed, both of these subregions peaked even earlier than M1. Such early peaks were observed in trials with relatively long RTs (Figure 3D), whereas the peak latency of the left M1 exhibited a minimal dependence on RT. Importantly, the lack of substantial differences at RT = 1 suggests that these early peaks are unlikely to be attributable solely to variability in the latency of the hemodynamic response.

We conducted a control experiment using a simple reaction task to test whether this early activation was attributable to motor output (Figure 3E). We calculated the mean BOLD signal within the 3- to 4-second window following the response for each subregion because the signals in the two subregions were noisy and did not yield clear peaks in this control experiment. When comparing between the Mooney task and the simple reaction task, the SC region exhibited significantly greater activation only in the Mooney task (Wilcoxon signed-rank test with Benjamini–Hochberg FDR correction^63^, q < 0.01), supporting its involvement beyond simple motor output. Among all 148 subregions, only the SC, bilateral fusiform gyrus, and cerebellar vermal lobules VIII-X showed significantly greater activation at 3–4 seconds after the response in the Mooney image task than in the simple reaction task (Extended Data Figure E5). In addition, the SC occupied a distinct location within the low-dimensional PCA space, separating it from other subregions in the C_RA_ (Figure 2B inset). Together, these results indicate that the SC is an early and functionally distinct contributor to neural dynamics for sudden insight.

### Directed dynamic interactions highlight the distinctive role of the superior colliculus

Although the SC exhibited an early peak in activity and emerged as a distinct contributor, further validation is necessary since the timing of the peak alone does not establish a causal link. We applied a data-driven dynamic systems approach based on empirical dynamic modeling (EDM) ^64–66^, which allows for the inference of dynamical causality among brain regions during recognition, to move beyond simple correlational observations and investigate causal relationships within a complex and interconnected neural system.

We first analyzed the directed dynamic relationships among six representative time series: the three functional clusters (C_SP_, C_SN_, and C_RA_), the SC subregion that peaked earlier than the response, the response-related subregion M1, and the stimulus on/off time series (Stim). This analysis was performed because the time series of the three major functional clusters, which were averaged across multiple subregions, provide more stable and reliable results than those of individual subregions. The results revealed multiple interpretable patterns aligned with known brain functions (Figure 4A(i) and Extended Data Figure E6A). The external stimulus input exerted a directional influence only on the C_SP_, a cluster containing visual areas, and appropriately, no neural signals were found to causally influence the stimulus itself. A mutual inhibitory relationship was identified between the C_SP_ and C_SN_, consistent with the well-established antagonistic interaction between the dorsal attention and default mode networks.

**Figure 4.**
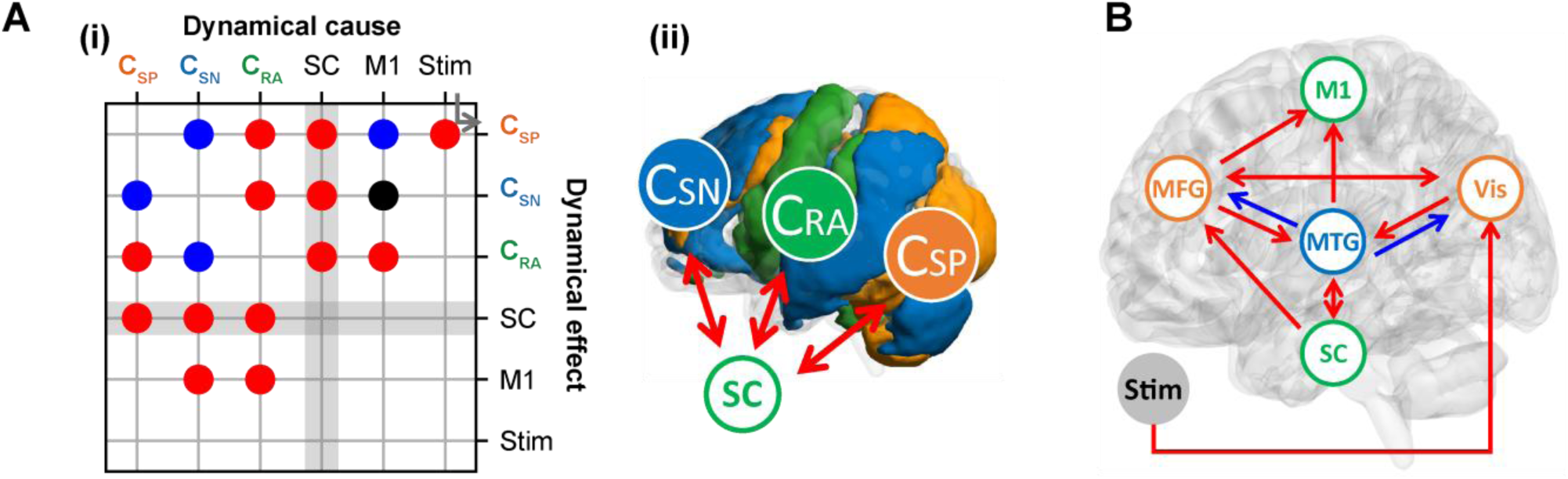
Directed dynamic interactions. **A**, Functional cluster-level analysis. (i) Interaction matrix. The blue, red, and black circles represent positive, negative, and mixed influences, respectively. C_SP_, C_SN_, and C_RA_ refer to the three functional clusters shown in Figure 2. The matrix is not symmetric across the diagonal because the relationships are directed, as exemplified by the gray arrow. The SC was the only region among all 148 subregions across the whole brain that exhibited mutual positive dynamic interactions with all three functional clusters (Extended Data Figure E6B), which represented whole-brain networks related to this insight task, as depicted in (ii). **B**, Schematic summary of interactions identified in the subregion-level analysis. Representative shortest paths from the stimulus to M1 that include the SC, selected from all pairwise interactions are shown. Vis represents visual areas such as the occipital gyrus, fusiform gyrus, and superior parietal lobule. The MTG (middle temporal gyrus) is representative of a group that includes the angular gyrus, lateral orbital gyrus, posterior cingulate gyrus, precuneus, supramarginal gyrus, and superior temporal gyrus. The MFG (middle frontal gyrus) represents regions including the opercular part of the inferior frontal gyrus and the supramarginal gyrus.

Importantly, we found that the SC was the only region among all 148 subregions across the whole brain that exhibited mutually positive dynamic interactions with all three functional clusters (Figure 4A(ii) and Extended Data Figure E6B). Despite the opposing behaviors of C_SP_ and C_SN_ in general, the SC exerted a positive causal influence on both, highlighting its unique role in mediating large-scale neural activation.

Furthermore, while M1, the motor cortex responsible for executing the button press, was influenced by both the C_RA_ and the C_SN_, it did not receive input from the SC. This dissociation suggests that the involvement of the SC extends beyond motor output and supports a more cognitive role in recognition-related processing.

In a more fine-grained analysis at the subregion level (Figure 4B and Extended Data Figure E6C), the shortest direct paths from the stimulus to M1 typically followed a sequential route through visual subregions and then through areas such as the middle temporal gyrus (MTG) or middle frontal gyrus (MFG), ultimately reaching M1. When considering the shortest paths from the stimulus to M1 that do include the SC, the SC was embedded within extended pathways that also passed through the MTG or both the MTG and the MFG. This pattern suggests that the SC may contribute to higher-order processing within large-scale cortico-subcortical networks, such as integrating visual and mnemonic, hypothesis generation and validation processes. This interpretation is consistent with our finding that the SC was among a few subregions that exhibited significantly greater activation during Mooney image recognition than during the simple reaction task.

## Discussion

In this study, we investigated neural dynamics during the open-ended recognition of Mooney images and reported two main findings: the characterization of three patterns in whole-brain neural dynamics and the identification of a distinctive role of the SC in those dynamics. Among these, the prominence of the SC is of particular importance. Recent research has documented the involvement of the superior colliculus (SC) in a wide range of tasks^41,44–54^, laying an important foundation for exploring its role in more complex cognitive functions. Building on this foundation, the present study provides compelling experimental evidence that the SC plays a critical role in an open-ended insight task, which may represent a particularly important function, as it supports adaptation to novel and challenging situations.

Open-ended insight tasks, such as those used in this study, are thought to involve a sudden, generative shift from unawareness to resolution, typically without intermediate stages^4,28–32^, which qualitatively differs from the gradual evidence accumulation seen in forced-choice paradigms. In the latter, the SC has been shown to implement decision thresholds^52,67^. Despite this qualitative distinction, the SC may serve a common computational function, i.e., threshold implementation, across both types of tasks. Indeed, our finding that the SC exhibits transient activation within a cortical network characterized by sustained activity (Figures 2 and 4) aligns well with observations in nonhuman primates^52^. As with studies on the SC in nonhuman primates^41,44–53^, however, our findings do not address the potential involvement of the SC in saccadic control^38,41,68^, unconscious visual processing^38–43^, or insight outside of the visual domain. These areas remain important topics for future research. Given that insight is a fundamental function for biological systems to cope with unexpected situations^6,69,70^ and that the SC is an evolutionarily conserved structure, it may play a comparable role across species. Future investigations into its contribution to open-ended insight tasks in both human and nonhuman models would be highly valuable.

In characterizing whole-brain neural dynamics, we found that each individual response pattern largely aligns with prior findings, yet our analysis integrates them into a unified spatiotemporal framework and reveals their coordinated temporal dynamics. For example, the clear separation between stimulus-driven activation in the visual cortex (C_SP_) and recognition-related activation in frontal areas (C_RA_) is consistent with earlier findings^25–27^. The DMN, which is largely captured in the C_SN_, showed characteristic task-related suppression, followed by a gradual increase in activity over time (clearly visible in Figure 2C(iii)), and finally transient activation at the moment of recognition. These patterns may reflect the known association of the DMN with creative cognition^4,71^.

A key contribution of this study is the identification of SC involvement using fMRI, despite its small size and deep location. In addition to detecting the SC and validating that its activity reflects a functionally prominent role, we showed that the SC is embedded within higher-order cortical interactions (Figure 4). EDM even detected an inhibitory influence from the substantia nigra to the SC (Extended Data Figure E6), reflecting a known local interaction within the midbrain^72,73^. While invasive approaches enable the direct interrogation of interactions between selected regions^61^, our use of EDM demonstrates that pairwise causal relationships across the entire brain can also be assessed noninvasively. This approach provides a powerful framework for future investigations of whole-brain network dynamics, including noncortical regions, and helps to advance a broader conceptual shift beyond cortex-centered models of brain function^74,75^. Moreover, EDM enabled stable and interpretable results based on long, continuous time series data, without relying on trial averaging. This approach holds promise for revealing mechanisms hidden in trial-by-trial neural dynamics that conventional averaging may have overlooked.

## Methods

### Participants

Fourteen healthy native Japanese speakers (seven females; mean age: 23.2 ± 2.5 (SD) years) participated in an fMRI experiment investigating Mooney image recognition. All the participants had normal or corrected-to-normal vision and successfully passed the visual object and space perception battery tests (Thames Valley Test Company, Bury St. Edmunds, Suffolk, England). All of these participants also participated in a separate fMRI experiment for the simple reaction time task, in which the participant pressed a button as quickly as possible when a white disk (0.6° visual angle in diameter) was briefly presented (for 0.2 s), overlapping the fixation point at the center of a black background. The interstimulus intervals were randomized, ranging from 3 to 20 seconds. This study was approved by the Ethics Committee for Human and Animal Research of the National Institute of Information and Communications Technology, Japan. All participants provided written informed consent in accordance with the committee’s guidelines and were compensated for their participation.

### Visual stimuli

We created novel Mooney images for this experiment using the following procedure. Ninety digital color images were selected from clip-art collections (Sozaijiten, Image Navi Corp., Japan) and personal photographs. Each image featured a distinct object that could be easily recognized to describe verbally (e.g., “a chicken’s head facing the right”). The selected images represented diverse categories to minimize categorical bias: human faces (5 images); humans with faces and bodies (20); natural or artificial scenes (6); hand-held tools (3); animals such as dogs, fish, and frogs (43); and vehicles (13). These color images were degraded using monochromatic binarization. The binarization threshold for each image was manually adjusted using Adobe Photoshop (Adobe Inc., USA) to ensure that the subjective difficulty of recognizing the degraded images was evenly distributed within an identifiable range. Presentation software (ver. 23.1, Neurobehavioral Systems) was used for stimulus presentation and response collection. The LCD monitor was positioned 184 cm from the participant’s eye level inside the MRI gantry. The stimuli were displayed at the center of the monitor, with each image measuring 640 pixels on its longer side, subtending a visual angle of 10° and featuring a white luminance of approximately 100 cd/m^2^ against a black background.

### Data acquisition

MRI data were acquired using a 3 T MR scanner (Vida, Siemens) at the Center for Information and Neural Networks. For functional imaging, a simultaneous multislice (SMS) sequence with six bands and ascending T2*-weighted gradient-echo echo-planar imaging (EPI) was used. This procedure produced transaxial slices covering the entire cerebrum and cerebellum, excluding the eyeballs, through oblique scanning. The imaging parameters were as follows: repetition time (TR) = 1000 ms, echo time (TE) = 30 ms, flip angle (FA) = 60°, matrix size = 100 ×100, number of slices = 72, and voxel size = 2.0 × 2.0 × 2.0 mm³. The scanner collected ten dummy volumes before EPI recording began to allow the stabilization of the functional imaging data. The acquired MRI data in raw DICOM format were converted to NIFTI format and preprocessed using Statistical Parametric Mapping 12 (SPM12^55,56^; University College London, United Kingdom; https://www.fil.ion.ucl.ac.uk/spm/), implemented in MATLAB 2020b (MathWorks). For each session, head motion-induced displacement in the scans was corrected through realignment relative to the mean image using affine transformation, with the image headers adjusted to reflect the relative displacement. Slice timing correction was applied to account for differences in signal acquisition times across slices, adjusting each time series as if all slices were acquired simultaneously with the reference slice. For SMS imaging with six bands, the reference slice was set at the middle of each scan band. Spatial normalization was performed using nonlinear three-dimensional transformations to align individual brain images with the ICBM template for East Asian brains, which was based on the MNI152 standard space. As a result of this normalization with resampling to 2.0 × 2.0 × 2.0 mm³ isotropic resolution, the functional images were transformed into a common space with dimensions of 79 × 95 × 79 voxels (totaling 592,895 voxels). The normalized images were subsequently smoothed using a three-dimensional Gaussian kernel with a full width at half maximum (FWHM) of 6 mm. The data from each session were high-pass filtered with a cutoff frequency of 0.01 Hz to eliminate low-frequency artifacts and task-unrelated trends. Subsequently, the data were standardized within each session to have a zero mean and unit variance.

### Experimental design

A challenge in fMRI experiments investigating Mooney image recognition is determining whether participants arrive at the correct solution without requiring verbal responses during MRI scanning. Verbal responses can introduce signal changes unrelated to Mooney image recognition itself in the MR environment. We leveraged a key empirical property of Mooney image recognition to address this issue: once the solution is known, a previously ambiguous image becomes immediately recognizable. Using this property, we developed a post hoc verification method in which each session consisted of the two parts described below. The first part of a session, referred to as the scanning part, aimed to acquire brain activity data related to Mooney image recognition and was conducted during MRI scanning. The second part, referred to as the verification part, was conducted immediately after the scanning part without MRI scanning, allowing participants to provide verbal reports to verify their answers post hoc. Details of the methods are described below.

Before data collection, the participants were provided with instructions about the task, which required them to observe each Mooney image with the aim of recognizing meaningful objects hidden within it and to press a button as soon as they identified a stable object. The instructions emphasized that only the button press was required during the scanning part, whereas verbal reporting was additionally required in the verification part. The participants practiced with several example images beforehand to become familiar with the procedure.

The experiment consisted of six sessions, each featuring 15 different Mooney images, which were commonly used in both the scanning and verification parts of the session. A single trial of the scanning part of a session began with a 15-second rest period with a white fixation point at the center of a black background (Figure 1A). This period was followed by a stimulus period, during which a Mooney image was presented. The order of the 15 images in each session was randomized for each participant. The participants were instructed to report their recognition moment by pressing an assigned button during the presentation of the Mooney image. The instructions allowed participants to press the button again if they identified another valid solution to prevent them from hesitating to respond. After the final response, the Mooney image remained on display for an additional 10 seconds before proceeding to the next trial. If a participant did not press the button at all during the 30-second presentation, the trial was classified as “no response,” indicating that the participant was unable to identify any solution. In such cases, the trial immediately transitioned to the next one.

In the verification part of each session, which was conducted immediately after the scanning part without MRI scanning, the same 15 Mooney images were presented in a different order. The participants were instructed to press a button as soon as they identified the object in the image. Upon pressing the button, the Mooney image was turned off, and the participants were asked to verbally report their recognition to the experimenter. If a participant’s answer was unclear, the experimenter asked follow-up questions to verify whether the object was correctly identified. A response in the scanning part was judged to be correct if the participant’s response in the verification part was both rapid (response time within 3.5 seconds) and validly matched the correct solution. For each participant, the 90 trials (90 different images) from the scanning parts across the six sessions were categorized into correct responses (i.e., matches with the corresponding original color image; 930 trials), incorrect responses (128 trials), or no responses (201 trials). The overall accuracy was high, which is consistent with prior findings that, in insight-based tasks, participants tend to either provide correct answers or omit responses, with incorrect responses being relatively rare ^31^. However, because many Mooney images are substantially degraded and often lack sufficient information to define a single objectively correct answer, our analysis focused on participants’ perceptual interpretations rather than strict correspondence with the original color images. Accordingly, all trials were included in subsequent analyses, regardless of the response accuracy. By carefully setting the difficulty of the images, we were able to obtain 153 trials in which participants took more than 8 seconds to recognize the Mooney images (Extended Data Figure E1). These trials included data from all 14 participants. The 8-second threshold corresponds to the time point at which the HRF returns to baseline, allowing us to dissociate recognition-related BOLD activity from stimulus-onset-related responses.

### Prescreening of task-associated regions

As a prescreening step, we excluded brain regions that did not show significant changes in BOLD activity during the recognition task to focus our subsequent analyses on regions potentially associated with Mooney image recognition. A general linear model (GLM) was constructed to identify brain regions exhibiting task-related activation. The model included three regressors corresponding to distinct temporal phases: preresponse (stimulus onset to response), response (a transient response event), and postresponse (response to stimulus offset). Each regressor was designed to capture both activations and deactivations relative to the baseline. This model was convolved with the canonical hemodynamic response function (HRF ^76^; the parameters were the same as those in the previous study ^77^) to generate the regressors. Although the canonical HRF was originally derived from activation-related BOLD responses, we applied the same HRF to model both activations and deactivations, as the GLM framework captures signal increases and decreases through the sign of the estimated beta weights. In the individual (first-level) analysis, preprocessed fMRI data for each participant, including realigned, slice-timing corrected, spatially normalized, smoothed, high-pass filtered, and standardized data, as described in the Data Acquisition section, were analyzed using the general linear model described above. Serial autocorrelation was modeled using a first-order autoregressive (AR(1)) approach. For the group-level (second-level) analysis, contrast estimates (beta values) obtained from each participant’s first-level GLM were entered into voxel-wise one-sample t tests across 14 participants ^78,79^. The resulting statistical map was thresholded using the false discovery rate (FDR) correction for multiple comparisons (Benjamini–Hochberg procedure ^63^, q < 0.01), without cluster-level correction methods based on random field theory, as the analysis included small brainstem structures such as the superior colliculus (31 voxels in volume).

We excluded voxels that did not fall within any labeled area of the Neuromorphometrics atlas provided in SPM12 to restrict our analysis to anatomically meaningful regions. While the Neuromorphometrics atlas provides MNI space volume data for 136 tissue-defined brain regions, we included 122 areas, excluding 14 structures lacking neuronal cell bodies (e.g., white matter) ^77^. Among the 130,707 voxels corresponding to these 122 atlas-defined regions, statistically significant effects were observed on 68,263 voxels, representing approximately half of the voxels. Each of the three temporal regressors (preresponse, response, and postresponse) was associated with significant activation in approximately 30,000 voxels, with substantial overlap observed among them. Furthermore, approximately half of the voxels that showed a positive beta value for one regressor overlapped with those showing a negative beta value for another regressor. These results and an overview of the anatomical distribution of these significantly positive and negative beta values are shown in Extended Data Figure E2. The spatial distribution of these voxels was widespread and dispersed, suggesting that a broad network of brain regions was involved in this task. These activations tended to form contiguous clusters rather than appearing as isolated points.

### Prefunctional parcellation and extraction of BOLD time courses for representative subregions

Given that functionally heterogeneous activity patterns can arise within a single atlas region, we applied a prefunctional parcellation procedure to further subdivide regions exhibiting internally divergent signal profiles. This parcellation step was performed independently of the GLM-based prescreening described above to maintain analytical flexibility.

Specifically, BOLD time series from all participants were concatenated (totaling 41,446 seconds), and pairwise distances between voxel time courses were computed within each atlas region. Hierarchical clustering using complete linkage was then applied to define functional subregions. Given that the voxel time series were standardized, the mean squared Euclidean distance between voxel pairs is mathematically proportional to 1−R, where R is the Pearson correlation coefficient and is widely used in functional connectivity analyses. A uniform clustering threshold was applied across all atlas regions, yielding a total of 300 functional subregions. We confirmed that increasing the resolution of this parcellation (e.g., to approximately 1,000 subregions) did not qualitatively affect the conclusions of this study.

Following the prefunctional parcellation, we defined the representative voxel for each subregion as the one whose time series exhibited the smallest Euclidean distance to the mean time series of all voxels within that subregion. This approach was adopted to reduce the influence of noisy or outlier voxels while preserving the central temporal characteristics of each subregion. We then retained only those subregions whose representative voxels fell within the GLM-based prescreened areas described above. Excluding a single subregion containing fewer than 10 voxels resulted in 148 subregions being selected for further analysis. These 148 subregions collectively encompassed approximately 64% of the total voxel volume within the 122 atlas-defined regions (Extended Data Figure E3). This spatial coverage was slightly greater than that of the original GLM-based prescreened regions because each subregion was selected based on the location of its representative voxel, which was within the prescreened areas, even though the subregion itself could extend into regions outside the GLM-based mask. Among these 148 subregions, the left precentral gyrus, which is expected to be involved in right-hand button presses, was preserved, whereas the corresponding region in the right hemisphere, which was not task related, was excluded, as expected.

For each of the 148 subregions, we computed a representative BOLD time series using a robust averaging approach. Rather than using a simple mean, which could be unduly influenced by noisy voxels distant from the central tendency, we selected the 10 voxels whose time series were closest (in Euclidean distance) to the subregion’s mean time series and averaged them. The resulting 148 representative time series, each spanning 41,446 seconds, were used in subsequent analyses.

### Trial averages for different reaction times

We aggregated all trials in which the response occurred after 18 seconds, under the assumption that these trials share similar BOLD time series both before and after the response (designated RT ≥ 18), to extract the features in long RT trials, e.g., greater than 18 seconds. The resulting averaged time series is shown in Figure 2A (N = 44). For example, in a trial of RT = 20, the measurement window from 18 to 19 seconds was trimmed and included in the average. The condition-specific average time series (e.g., for RT = 8) and the aggregated average time series across trials with RT ≥ 8 are shown in Extended Data Figure E4.

### Large-scale functional clustering and model fitting

We first performed a principal component analysis (PCA) on the full time series data (148 subregions × 41,446 time points) to reduce the temporal dimensionality, which yielded 148 temporally compressed features (148 subregions × 148 temporal features). We then applied Gaussian mixture modeling using all 148 temporal features to perform clustering across a range of cluster numbers using scikit-learn ^80^. Figure 2B shows the Bayesian information criterion (BIC) scores for each model, indicating that a three-cluster solution provides the most efficient representation. The log-likelihood of the three-cluster model reached 62% of the difference between the 1-cluster model and the full-dimensional model, indicating the efficiency of this solution. Although the clustering was performed in 148 PCA dimensions, visualizing the data in the PCA space of the first two components suggested a reasonable separation into three groups (inset of Figure 2B).

We then fitted the time series of the three functional clusters using a model composed of four event-related components: stimulus onset, response, stimulus offset, and sustained stimulus presentation (Figure 2C(i)). Each component was convolved with the canonical hemodynamic response function (HRF), and the model was fitted separately to the average time course of each functional cluster, with 12 parameters in total (i.e., four components for each of the three clusters), using SciPy ^81^.

### Detection of the peak time just after the response

We analyzed trials with an RT of 8 seconds or longer (N = 153) to estimate the timing of peak activation following the response. For each trial, we extracted BOLD signal data from a 12-second window beginning 2 seconds before the response. Because peak detection at the single-trial level is highly susceptible to noise, a moving average was applied with a 2-second window before and after each time point, and we employed a nonparametric bootstrap approach to improve robustness. Specifically, 153 trials were resampled with replacement 1000 times to generate averaged time series, from which the peak time was computed for each iteration. This procedure yielded a bootstrap distribution of peak timings, from which we derived the mean, standard deviation, and 95% confidence interval. This resampling-based estimation is advantageous for capturing central tendencies and variability without relying on parametric assumptions.

We ensured that only subregions with clear and temporally localized peaks were included by excluding those in which any of the 1000 bootstrap samples showed a peak at either edge of the 12-second window. We also excluded subregions in which the maximum signal within the window did not exceed both endpoints by at least 0.1. Among the remaining subregions, those whose 95% confidence intervals were entirely within at most 5 seconds after the response are shown in Figure 3A. Figure 3D shows the same analysis performed across a range of RT bins.

### Empirical dynamic modeling

We performed empirical dynamic modeling using the pyEDM package^82^, which was developed and maintained by the original creators of the method ^64^. Using established analytical procedures^64,65^, we assessed the presence of dynamic relationships using convergent cross mapping (CCM). We then applied the multivariate sequential locally weighted global linear map (S-map) method to assess the direction (positive or negative) of the influence for each detected dynamic relationship. A brief description of this analysis, together with the data and detailed methods, is provided in the Extended Data section (Extended Data Figure E6).

Analyses were conducted in accordance with standard documentation^82^, with minor adjustments to two parameters, “E” and “theta.” While these parameters are typically optimized for each time series, their estimation is often unstable, depending on the library (time window) used. We observed similar variability in our data; however, the optimal values consistently fell within the range of 5 to 10 for E and 0.3 to 2 for theta. In particular, for the three functional clusters, which showed particularly stable dynamics, 6 for E and 1 for theta were most commonly observed. Therefore, we used these fixed values for all the time series. Note that this choice was further justified by the fact that all time series were derived from BOLD signals and thus expected to share similar nonlinear temporal characteristics within the 5 to 10 second range. Additionally, the stimulus time series lacks a meaningful embedding structure, and a fixed embedding dimension ensures analytical consistency with the other BOLD-based signals.

## Data availability

The fundamental data generated in this study (time course and brain map data of all subregions and three functional clusters, comprising 41,446 time points from 14 human participants) are available on Zenodo and will be made publicly available upon publication. During peer review, the data can be accessed via the following private link: [Private link for reviewers only; not intended for public dissemination prior to formal publication].

## Code availability

The code supporting the findings of this study is available from the same Zenodo repository and will also be made publicly accessible upon publication. During peer review, the code can be accessed via the same private link as the data.

## Acknowledgments

We are grateful to Drs. Shigeru Kitazawa, Tetsuya Shimokawa, Atsushi Yokoi, and Ben Seymour for their invaluable input, and the instrumental operators, cluster computing teams, and assistants for their expert support. This study was supported in part by MIC under a grant entitled “R&D of ICT Priority Technology (JPMI00316)” and JSPS KAKENHI Grant Numbers JP24K15188 and JP25H01365.

## Author contributions

T.M., T.Y., and K.H. designed the study and developed the concept. I.O., K. Kaneko, and T.Y. provided supervision for the study. T.M. and K. Kunishige collected the data. S.S. and K.H. analyzed the data. I.O., K. Kaneko, and M.H. provided insights regarding data interpretation. M.H. and K.H. drafted the manuscript. All the authors contributed critical revisions and approved the final manuscript for publication.

## Additional information

Correspondence and requests for materials should be addressed to Kazufumi Hosoda. Reprints and permissions information is available at www.nature.com/reprints.

## Extended data

**Extended Data Figure E1.**
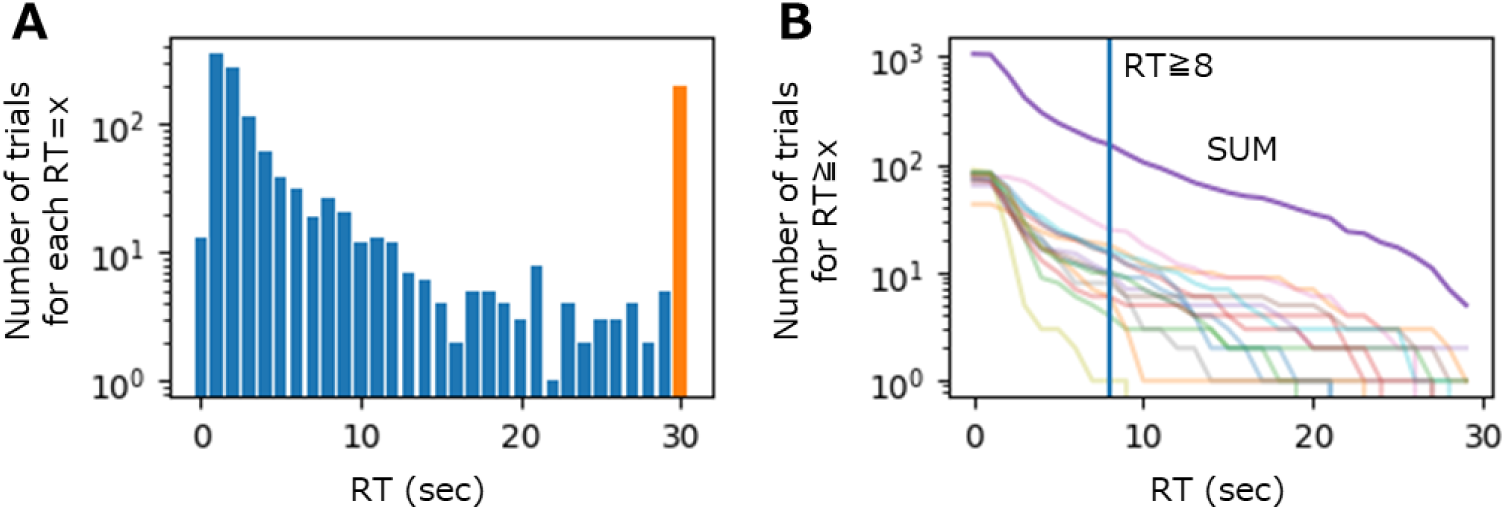
Distribution of reaction times (RTs) across trials. A. RT distribution across all participants. Here, for example, RT = 1 refers to trials where the response occurred between 1 and 2 seconds after stimulus onset. The bar at RT = 30 represents the frequency of no-response trials. B. Cumulative RT distribution per participant. For each participant, the number of trials with an RT equal to or longer than the value on the x-axis is plotted. While one participant responded markedly faster than the others did, the rest showed no notable bias. All participants contributed to trials with RT ≥ 8, and all but one contributed to trials with RT ≥ 18. Although longer RT trials were less frequent overall, the appropriate stimulus difficulty allowed us to obtain 153 trials with an RT ≥ 8 seconds.

**Extended Data Figure E2.**
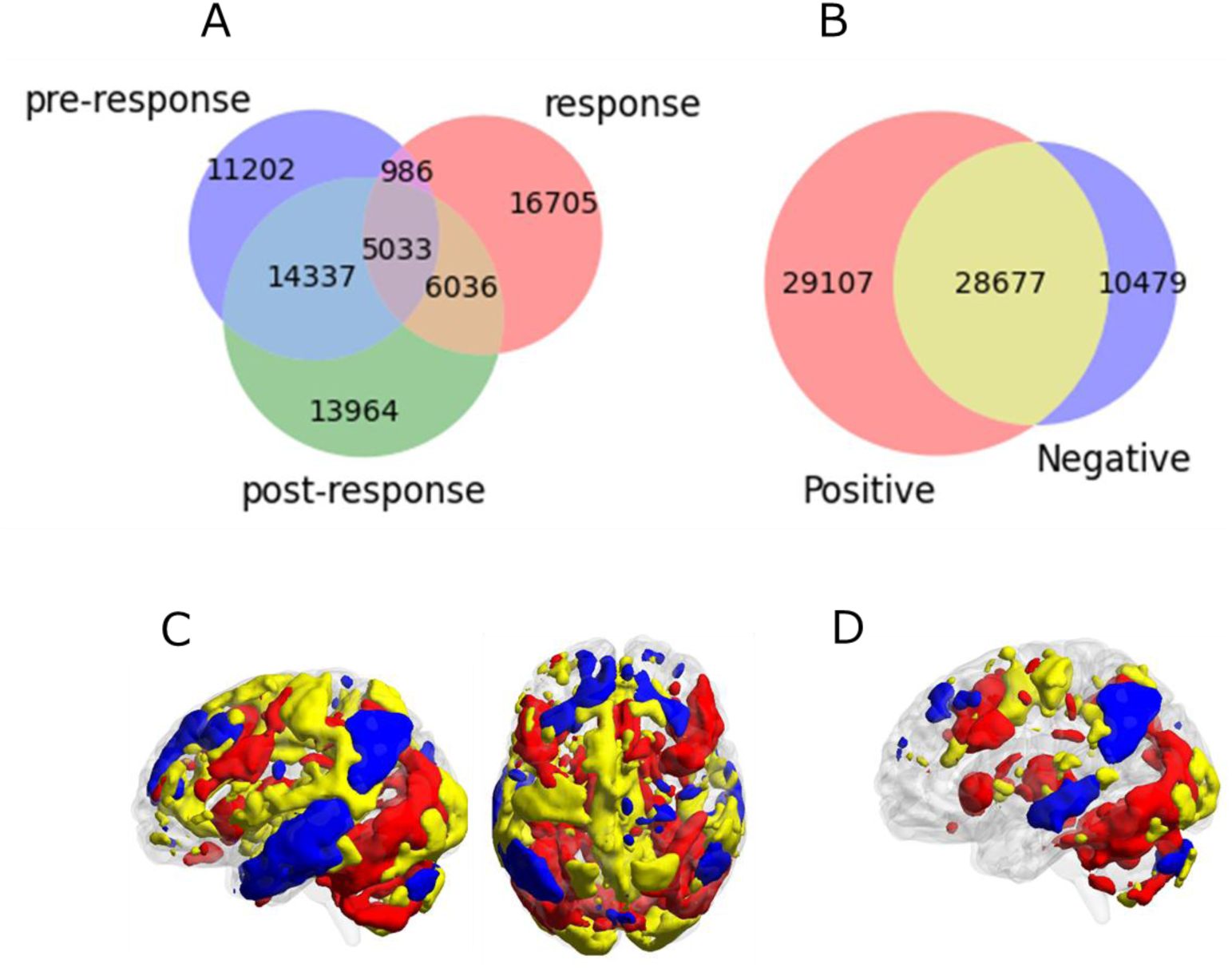
Prescreening of task-associated regions. **A**, Venn diagram showing the number of voxels significantly associated with each of the three distinct temporal phases: preresponse (stimulus onset to response), response (a transient event at response), and postresponse (response to stimulus offset). Each of the three temporal regressors was associated with significant activation in approximately 30,000 voxels, with substantial overlap observed among them. **B**, Venn diagram illustrating the overlap of voxels that showed significant effects for at least one of the three regressors, classified by whether they exhibited a positive or negative beta value. **C**, Anatomical distribution of the voxel clusters depicted in Panel B, color-coded according to beta polarity. **D**, Spatial distribution of voxels that survived a more stringent multiple-comparisons correction using the Benjamini–Yekutieli method (shown as an illustrative example). Even when such a more stringent multiple-comparisons correction was applied, the overall conclusions remained unchanged.

**Extended Data Figure E3.**
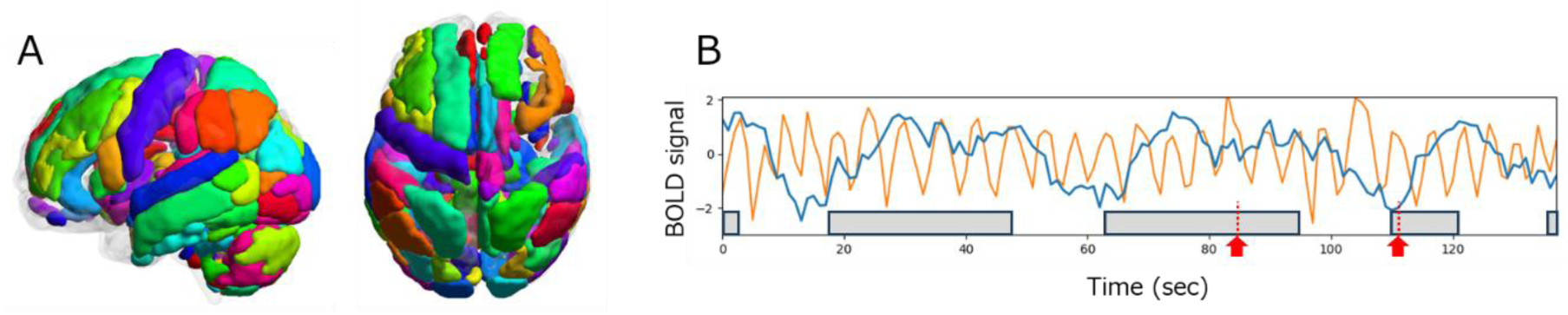
Prefunctional parcellation and extraction of BOLD time courses for representative subregions. **A**, A total of 148 subregions identified via prefunctional parcellation and retained based on the GLM-based prescreening. Each color represents a different subregion; colors are used for visualization purposes only and do not carry specific meanings. For example, the left precentral gyrus, which is expected to be involved in right-hand button presses, was preserved (purple), whereas the corresponding region in the right hemisphere, which was not task related, was excluded, as expected. **B**, Representative BOLD time series from a subregion included in the prescreening (blue) and from one excluded by the prescreening (orange). The gray boxes indicate stimulus presentation periods, and the red arrows mark the time points of response. The blue trace appeared to indicate stimulus-related responses. In contrast, the orange trace displayed oscillatory fluctuations that appeared unrelated to the stimulus presentation.

**Extended Data Figure E4.**
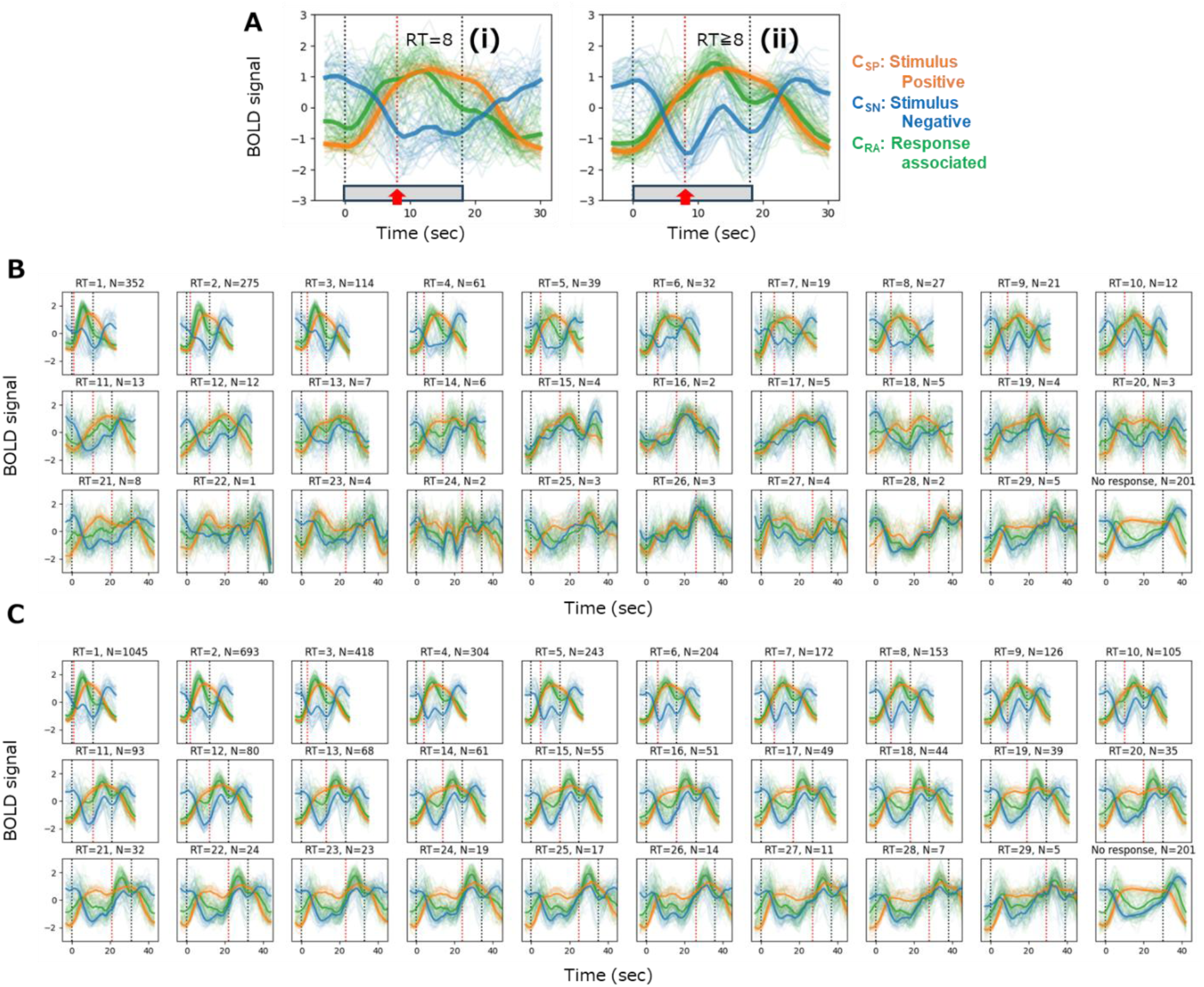
Trial averages for different reaction times. A. Representative condition-specific average time series (RT = 8, (i)) and the aggregated average time series (RT ≥ 8, (ii)). Thin lines represent individual subregions, color-coded by functional cluster identity (orange: C_SP_; blue: C_SN_; green: C_RA_). The thick lines represent the average time course within each functional cluster. The gray boxes between the two black dotted lines indicate the periods during which the stimuli were displayed, and the red arrow with the red dotted line indicates the moment of the button press response. **B**. Condition-specific average time series. **C**. Aggregated average time series. The two black dotted lines and the red dotted line have the same meanings as those in Panel A.

**Extended Data Figure E5.**
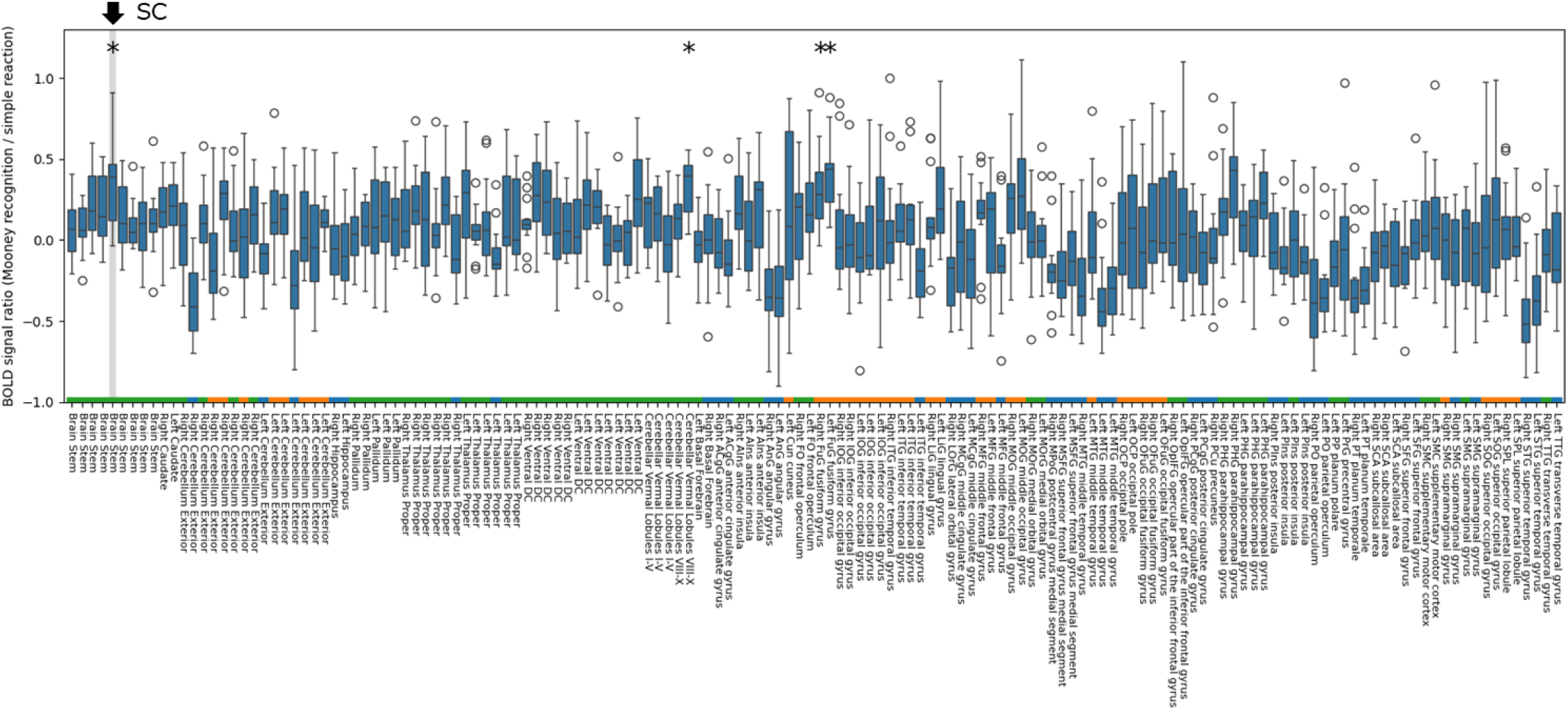
Comparison of BOLD signal amplitudes between the Mooney image recognition task and the simple reaction task across all 148 subregions. The three colors preceding each label indicate the functional cluster membership of the corresponding subregion. Among all 148 subregions, only the SC (“Brain Stem,” indicated by black arrow), bilateral fusiform gyrus, and cerebellar vermal lobules VIII-X showed significantly greater activation in the Mooney image task than in the simple reaction task (asterisks; Wilcoxon signed-rank test with Benjamini–Hochberg FDR correction, q < 0.01).

**Extended Data Figure E6.**
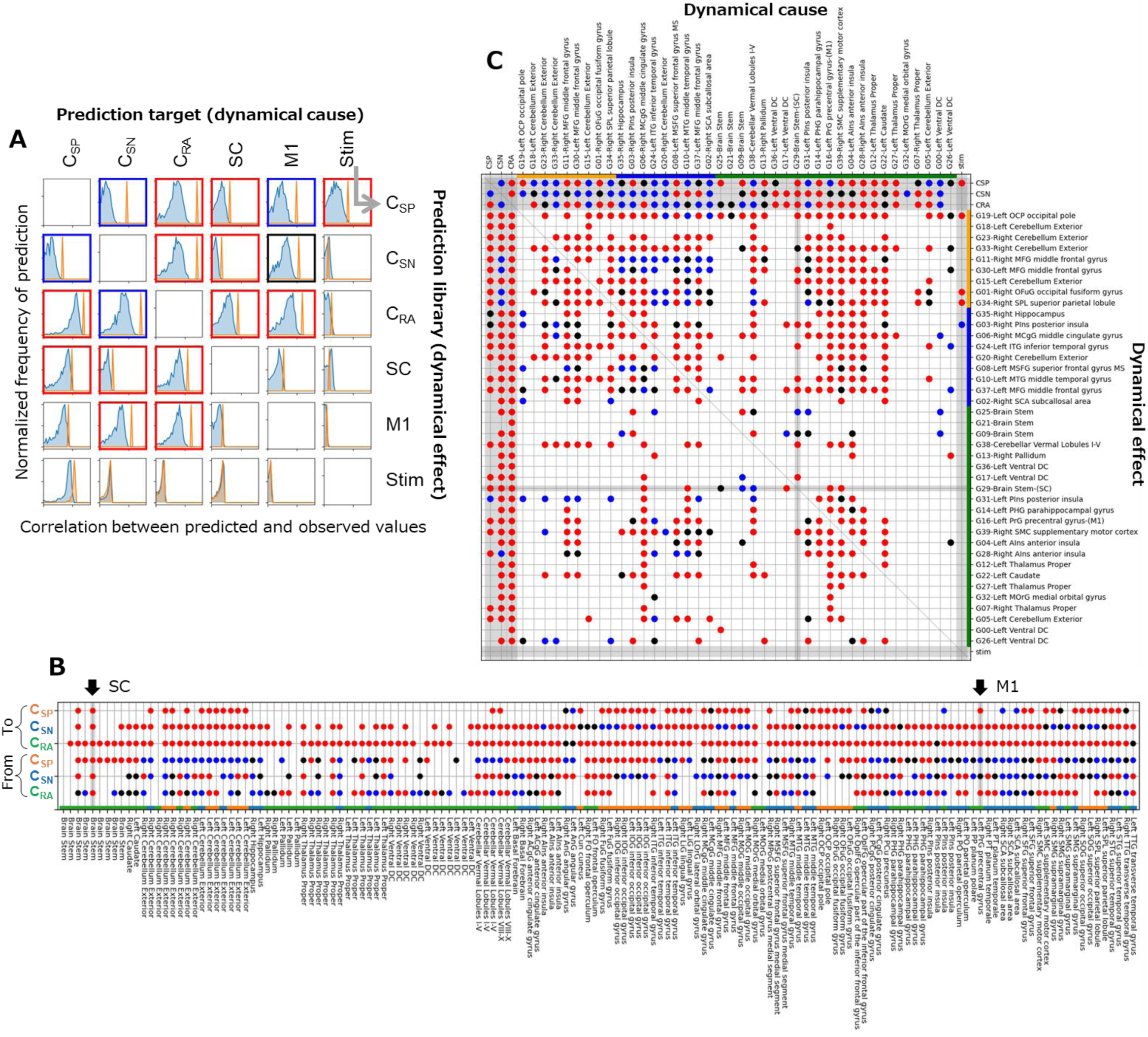
Empirical dynamic modeling (EDM).

We begin with a brief description of this analysis. First, we assessed the presence of dynamic relationships using convergent cross mapping (CCM) ^64^. As an example illustrating this model, let us consider the relationship between the C_SP_ and the stimulus time series. Expecting that activity in the C_SP_, which includes visual areas, would be influenced by the stimulus is reasonable. We attempted to predict the stimulus time series using the C_SP_ time series to test this assumption. If the prediction accuracy when the full-length C_SP_ time series is used is significantly greater than that when shorter segments are used, the C_SP_ time series data can be interpreted as containing embedded information about the stimulus. In dynamic systems, this result corresponds to the presence of a stimulus-related term in the differential equation governing C_SP_ (i.e., *dC*/*dt* = *f*(*C*, *S*, …), where *C* and *S* are the states of the C_SP_ and stimulus, respectively), implying that the stimulus exerts a dynamic influence on the C_SP_. Importantly, in this framework, the prediction target represents the dynamic cause, which is the inverse of the conventional terminology. Conversely, predicting C_SP_ from the stimulus may also yield above-chance accuracy, as C_SP_ is influenced by the stimulus; however, since C_SP_ does not influence the stimulus variation, the stimulus time series does not embed information about C_SP_, and the prediction accuracy will not improve with increasing data length (library size). The pattern in which the prediction accuracy increases and asymptotically converges as more data are used is referred to as convergence, and this pattern is used as a criterion for detecting dynamical causality in CCM. The prediction accuracy was evaluated by calculating Spearman’s rank correlation coefficient between the predicted and observed signals ^64–66^. Specifically, for the analysis focusing on the relationship with the three functional clusters (Panels A, B, and Figure 4A), we defined the presence of a dynamic relationship when the lower bound of the 99% CI for the prediction accuracy using the maximum library size (i.e., the full time series of 41,446 data points) exceeded the upper bound of the 99% CI using the minimum library size (8 data points). Each CI was estimated from 1,000 predictions. For the subregion-level analysis (Panel C and Figure 4B), we used a 95% CI instead of a 99% CI, as detecting subregion-wise interactions is inherently more difficult than detecting interactions at the cluster level, which is averaged across multiple subregions. We assessed the direction (positive or negative) of the influence for each detected dynamic relationship by applying the multivariate sequential locally weighted global linear map (S-map) method ^64,65^. This approach estimates, e.g., the derivative ∂*C*/∂*S*, which represents the change in C_SP_ in response to a change in the stimulus, based on their respective time series. The sign of this derivative indicates whether the influence is excitatory (positive) or inhibitory (negative). Because the derivative is calculated at each time point, we summarize the sign of interaction (I_S_) using a standardized index defined as I_S_ = M/SD, where M and SD are the mean and standard deviation of the derivative values, respectively. We defined positive and negative influences when I_S_ ≥ 2 and I_S_ ≤ −2, respectively (red and blue marked). We defined a mixed influence when - 2 < I_S_ < 2 because the influence fluctuates between positive and negative in this case.

We used only the first of six sessions from each participant in all analyses to reduce the computational demands. Before making this decision, we confirmed in the cluster-level analysis that results based on all six sessions and those based only on the first session produced similar outcomes, with the cluster-level analysis providing sufficient stability for comparison. This approach reduced the total computational load to approximately one-hundredth of that required for full-session analyses.

### A. Dynamic relationships in the cluster-level analysis

Each subplot shows histograms of the prediction accuracy for one prediction library to the target pair. The orange histogram corresponds to predictions using the maximum library size (entire time series), whereas the blue histogram reflects predictions using the minimum library size (8 continuous time points). The subplots outlined with bold borders indicate statistically significant dynamic relationships. The color of the border represents the sign of the interaction: red indicates a consistently positive influence, blue indicates a consistently negative influence, and black indicates mixed polarity, where the direction of influence varied over time. These results are summarized in Figure 4A.

### B. Interactions between each subregion and each of the three functional clusters

Directed dynamic interactions were estimated using the time series from all 148 subregions and the representative time series of the three functional clusters. The three colors preceding each label indicate the functional cluster membership of the corresponding subregion. Among all subregions, only the SC exhibited bidirectional positive interactions with all three clusters. The result for the interaction from C_SP_ to M1 differs from Panel A, despite the use of identical analytical procedures. This discrepancy reflects a borderline case with respect to the 99% CI and is included here as such a representative example. Importantly, this difference has no bearing on the main conclusions of the study. We confirmed that such marginally unstable results were not used in formulating the main conclusions of the study.

### C. Subregion-level interactions

We first selected 40 representative subregions from the original set of 148 to reduce the computational complexity of the pairwise interaction analysis. The original subregions were based on parcellations within SPM12-defined atlas regions, which often include anatomically similar areas as separate entries, such as left and right homologs. Analyzing all pairwise interactions among the 148 subregions would result in an excessive number of comparisons, and thus we excluded those subregions with highly similar time series and retained a smaller representative subset. This approach reduced the total computational load to approximately one-tenth of that required for full subregion-level analyses.

For this selection, we applied principal component analysis (PCA) for dimensionality reduction, followed by Gaussian mixed modeling (GMM), as in the main clustering analysis. We use the term “group” here instead of “cluster” to avoid confusion with the previously defined clusters. Specifically, we used the top 10% of PCA components (15 dimensions) to capture the main variance and selected 40 groups. This number was chosen because the log-likelihood of the 40-group model reached 97% of the difference between the 1-group model and the 148-group model, exceeding that of the 30-group model by more than 1% but differing by less than 1% from that of the 50-group model.

For each group, we computed a weighted average time series from the voxels of the subregions it contained and designated the subregion closest in time series distance to this group average as its representative. We did not use the weighted average time series for each group to avoid introducing bias across groups due to differences in the effects of averaging. Instead, we used the time series of the representative subregion directly as the representative signal for its group. Therefore, the grouping into 40 subregions served solely to exclude the remaining 108 subregions. Nonetheless, we retained information about which subregions were associated with each representative subregion. As a result, the SC subregion remained as a single-member group, whereas the M1 subregion, which included 2,557 voxels, was grouped together with eight other subregions of only a few hundred voxels each but was selected as the representative subregion.

This panel shows the estimated bidirectional dynamic interactions among all 44 elements: the 40 representative subregions, the three functional clusters, and the stimulus. The meaning of the circle colors follows the same convention as in Panels A and B and Figure 4A. The three colors preceding each label indicate the functional cluster membership of the corresponding subregion. Although only 40 subregions were analyzed in this context, each group often contained multiple anatomically related regions. For example, the group represented by OFuG (G01) included the left and right inferior occipital gyri, middle occipital gyri, lingual gyri, occipital fusiform gyri, and superior occipital gyri.

In this subregion-level analysis, as noted above, we applied a 95% confidence threshold for CCM, which differs from the thresholds used in Panels A and B. Nevertheless, the main findings remained unchanged: no brain regions were found to exert a direct influence on the stimulus; only the SC exhibited mutual positive interactions with all three functional clusters; no direct influence was detected from the SC to M1; and no direct influence was observed from the stimulus to the SC. These consistent results support the robustness of the main conclusions presented in the main text.

These results reveal more detailed pathways of information transmission than those identified in the cluster-level analysis. For example, if we search for the shortest path composed solely of positive interactions from the stimulus to M1 (G16), one possible route proceeds through OFuG (G01), then MTG (G10), and finally reaches M1. Several of these three-step paths exist, none of which include the SC. In contrast, in paths that do include the SC, such as a loop from the MTG to the SC and back, or from the MTG through the SC and then through the MFG (G30 or G37) before reaching M1, the SC is located in the middle of five-step routes. These representative routes are summarized in Figure 4B.

Other expected interactions were also reliably detected. For example, even within the small midbrain region, an inhibitory effect of the substantia nigra (SNr, included within the region labeled G09) on the SC was identified, reflecting a known local interaction^72,73^.

## References

1 Kaplan, C. A. & Simon, H. A. In search of insight. Cognitive psychology 22, 374–419 (1990).

2 Sternberg, R. J. & Davidson, J. E. The nature of insight. (The MIT Press, 1995).

3 Ahissar, M. & Hochstein, S. Task difficulty and the specificity of perceptual learning. Nature 387, 401–406 (1997). 10.1038/387401a0

4 Kounios, J. & Beeman, M. The cognitive neuroscience of insight. Annual review of psychology 65, 71–93 (2014).

5 Sprugnoli, G. et al. Neural correlates of Eureka moment. Intelligence 62, 99–118 (2017). 10.1016/j.intell.2017.03.004

6 Barabási, D. L., Ferreira Castro, A. & Engert, F. Three systems of circuit formation: assembly, updating and tuning. Nature Reviews Neuroscience 26, 232–243 (2025). 10.1038/s41583-025-00910-9

7 Mooney, C. M. Age in the development of closure ability in children. Can J Psychol 11, 219–226 (1957). 10.1037/h0083717

8 Dolan, R. J. et al. How the brain learns to see objects and faces in an impoverished context. Nature 389, 596–599 (1997). 10.1038/39309

9 Tallon-Baudry, C. & Bertrand, O. Oscillatory gamma activity in humans and its role in object representation. Trends in cognitive sciences 3, 151–162 (1999).

10 McKeeff, T. J. & Tong, F. The timing of perceptual decisions for ambiguous face stimuli in the human ventral visual cortex. Cerebral Cortex 17, 669–678 (2007).

11 Giovannelli, F. et al. Involvement of the parietal cortex in perceptual learning (Eureka effect): An interference approach using rTMS. Neuropsychologia 48, 1807–1812 (2010). 10.1016/j.neuropsychologia.2010.02.031

12 Castelhano, J., Rebola, J., Leitao, B., Rodriguez, E. & Castelo-Branco, M. To perceive or not perceive: the role of gamma-band activity in signaling object percepts. PloS one 8, e66363 (2013).

13 Kizilirmak, J. M., Galvao Gomes da Silva, J., Imamoglu, F. & Richardson-Klavehn, A. Generation and the subjective feeling of “aha!” are independently related to learning from insight. Psychological Research 80, 1059–1074 (2016). 10.1007/s00426-015-0697-2

14 Hsieh, P.-J., Vul, E. & Kanwisher, N. Recognition alters the spatial pattern of FMRI activation in early retinotopic cortex. Journal of neurophysiology 103, 1501–1507 (2010).

15 Gorlin, S. et al. Imaging prior information in the brain. Proceedings of the National Academy of Sciences 109, 7935–7940 (2012). doi:10.1073/pnas.1111224109

16 Hegdé, J. & Kersten, D. A link between visual disambiguation and visual memory. Journal of Neuroscience 30, 15124–15133 (2010).

17 Teufel, C. et al. Shift toward prior knowledge confers a perceptual advantage in early psychosis and psychosis-prone healthy individuals. Proc Natl Acad Sci U S A 112, 13401–13406 (2015). 10.1073/pnas.1503916112

18 Chang, R., Baria, A. T., Flounders, M. W. & He, B. J. Unconsciously elicited perceptual prior. Neuroscience of Consciousness 2016 (2016). 10.1093/nc/niw008

19 Martinsen, M. M. et al. Facial ambiguity and perception: How face-likeness affects breaking time in continuous flash suppression. J Vis 24, 18 (2024). 10.1167/jov.24.9.18

20 Jiang, Y. & He, S. Cortical Responses to Invisible Faces: Dissociating Subsystems for Facial-Information Processing. Current Biology 16, 2023–2029 (2006). 10.1016/j.cub.2006.08.084

21 Andrews, T. J. & Schluppeck, D. Neural responses to Mooney images reveal a modular representation of faces in human visual cortex. NeuroImage 21, 91–98 (2004). 10.1016/j.neuroimage.2003.08.023

22 Lu, Y. & Singer, W. Dynamic signatures of the Eureka effect: an EEG study. Cerebral Cortex 33, 8679–8692 (2023). 10.1093/cercor/bhad150

23 González-García, C., Flounders, M. W., Chang, R., Baria, A. T. & He, B. J. Content-specific activity in frontoparietal and default-mode networks during prior-guided visual perception. eLife 7, e36068 (2018). 10.7554/eLife.36068

24 Flounders, M. W., González-García, C., Hardstone, R. & He, B. J. Neural dynamics of visual ambiguity resolution by perceptual prior. Elife 8 (2019). 10.7554/eLife.41861

25 Imamoglu, F., Kahnt, T., Koch, C. & Haynes, J. D. Changes in functional connectivity support conscious object recognition. Neuroimage 63, 1909–1917 (2012). 10.1016/j.neuroimage.2012.07.056

26 Imamoglu, F., Koch, C. & Haynes, J.-D. MoonBase: Generating a database of two-tone Mooney images. Journal of Vision 13, 50–50 (2013).

27 Casile, A. et al. Neural correlates of minimal recognizable configurations in the human brain. Cell Reports 44, 115429 (2025). 10.1016/j.celrep.2025.115429

28 Murata, T., Hamada, T., Shimokawa, T., Tanifuji, M. & Yanagida, T. Stochastic Process Underlying Emergent Recognition of Visual Objects Hidden in Degraded Images. PLOS ONE 9, e115658 (2014). 10.1371/journal.pone.0115658

29. Hosoda, K., Seno, S. & Murata, T. Simulating Reaction Time for Eureka Effect in Visual Object Recognition Using Artificial Neural Network. IIAI Letters on Informatics and Interdisciplinary Research 3 (2023).

30. Smith, R. W. & Kounios, J. Sudden insight: All-or-none processing revealed by speed–accuracy decomposition. Journal of Experimental Psychology: Learning, Memory, and Cognition 22, 1443 (1996).

31 Kounios, J. et al. The origins of insight in resting-state brain activity. Neuropsychologia 46, 281–291 (2008). 10.1016/j.neuropsychologia.2007.07.013

32 Salvi, C., Bricolo, E., Kounios, J., Bowden, E. & Beeman, M. Insight solutions are correct more often than analytic solutions. Think Reason 22, 443–460 (2016). 10.1080/13546783.2016.1141798

33 Ludmer, R., Dudai, Y. & Rubin, N. Uncovering camouflage: amygdala activation predicts long-term memory of induced perceptual insight. Neuron 69, 1002–1014 (2011). 10.1016/j.neuron.2011.02.013

34 Becker, M., Yu, Y. & Cabeza, R. The influence of insight on risky decision making and nucleus accumbens activation. Sci Rep 13, 17159 (2023). 10.1038/s41598-023-44293-2

35 Tik, M. et al. Ultra-high-field fMRI insights on insight: Neural correlates of the Aha!-moment. Hum Brain Mapp 39, 3241–3252 (2018). 10.1002/hbm.24073

36 Ogawa, T., Aihara, T., Shimokawa, T. & Yamashita, O. Large-scale brain network associated with creative insight: combined voxel-based morphometry and resting-state functional connectivity analyses. Scientific Reports 8, 6477 (2018). 10.1038/s41598-018-24981-0

37 Goold, J. E. & Meng, M. Visual Search of Mooney Faces. Front Psychol 7, 155 (2016). 10.3389/fpsyg.2016.00155

38 Horwitz, G. D. & Newsome, W. T. Separate Signals for Target Selection and Movement Specification in the Superior Colliculus. Science 284, 1158–1161 (1999). doi:10.1126/science.284.5417.1158

39 Tamietto, M. & de Gelder, B. Neural bases of the non-conscious perception of emotional signals. Nature Reviews Neuroscience 11, 697–709 (2010). 10.1038/nrn2889

40 Ajina, S., Pollard, M. & Bridge, H. The Superior Colliculus and Amygdala Support Evaluation of Face Trait in Blindsight. Front Neurol 11, 769 (2020). 10.3389/fneur.2020.00769

41 Benarroch, E. What Are the Functions of the Superior Colliculus and Its Involvement in Neurologic Disorders? Neurology 100, 784–790 (2023). 10.1212/wnl.0000000000207254

42 Le, Q. V. et al. A Prototypical Template for Rapid Face Detection Is Embedded in the Monkey Superior Colliculus. Front Syst Neurosci 14, 5 (2020). 10.3389/fnsys.2020.00005

43 Lim, C., Inagaki, M., Shinozaki, T. & Fujita, I. Analysis of convolutional neural networks reveals the computational properties essential for subcortical processing of facial expression. Scientific Reports 13, 10908 (2023). 10.1038/s41598-023-37995-0

44 Wang, L., McAlonan, K., Goldstein, S., Gerfen, C. R. & Krauzlis, R. J. A Causal Role for Mouse Superior Colliculus in Visual Perceptual Decision-Making. The Journal of Neuroscience 40, 3768–3782 (2020). 10.1523/jneurosci.2642-19.2020

45 Krauzlis, R. J., Lovejoy, L. P. & Zénon, A. Superior colliculus and visual spatial attention. Annu Rev Neurosci 36, 165–182 (2013). 10.1146/annurev-neuro-062012-170249

46 Rahmati, M., DeSimone, K., Curtis, C. E. & Sreenivasan, K. K. Spatially Specific Working Memory Activity in the Human Superior Colliculus. J Neurosci 40, 9487–9495 (2020). 10.1523/jneurosci.2016-20.2020

47 Basso, M. A., Bickford, M. E. & Cang, J. Unraveling circuits of visual perception and cognition through the superior colliculus. Neuron 109, 918–937 (2021). 10.1016/j.neuron.2021.01.013

48 Thomas, A. et al. Superior colliculus bidirectionally modulates choice activity in frontal cortex. Nat Commun 14, 7358 (2023). 10.1038/s41467-023-43252-9

49 Lo, C.-C. & Wang, X.-J. Cortico–basal ganglia circuit mechanism for a decision threshold in reaction time tasks. Nature Neuroscience 9, 956–963 (2006). 10.1038/nn1722

50 Crapse, T. B., Lau, H. & Basso, M. A. A Role for the Superior Colliculus in Decision Criteria. Neuron 97, 181–194.e186 (2018). 10.1016/j.neuron.2017.12.006

51 Jun, E. J. et al. Causal role for the primate superior colliculus in the computation of evidence for perceptual decisions. Nat Neurosci 24, 1121–1131 (2021). 10.1038/s41593-021-00878-6

52 Stine, G. M., Trautmann, E. M., Jeurissen, D. & Shadlen, M. N. A neural mechanism for terminating decisions. Neuron 111, 2601–2613.e2605 (2023). 10.1016/j.neuron.2023.05.028

53 Peysakhovich, B. et al. Primate superior colliculus is causally engaged in abstract higher-order cognition. Nature Neuroscience 27, 1999–2008 (2024). 10.1038/s41593-024-01744-x

54 Schröder, S. et al. Arousal Modulates Retinal Output. Neuron 107, 487–495.e489 (2020). 10.1016/j.neuron.2020.04.026

55 Penny, W. D., Friston, K. J., Ashburner, J. T., Kiebel, S. J. & Nichols, T. E. Statistical parametric mapping: the analysis of functional brain images. (Elsevier, 2011).

56 Friston, K. J. et al. Statistical parametric maps in functional imaging: a general linear approach. Human brain mapping 2, 189–210 (1994).

57 Whitfield-Gabrieli, S. & Nieto-Castanon, A. Conn: a functional connectivity toolbox for correlated and anticorrelated brain networks. Brain connectivity 2, 125–141 (2012).

58 Xia, M., Wang, J. & He, Y. BrainNet Viewer: a network visualization tool for human brain connectomics. PloS one 8, e68910 (2013).

59 Buckner, R. L., Krienen, F. M., Castellanos, A., Diaz, J. C. & Yeo, B. T. The organization of the human cerebellum estimated by intrinsic functional connectivity. J Neurophysiol 106, 2322–2345 (2011). 10.1152/jn.00339.2011

60 Cunnington, R., Windischberger, C., Deecke, L. & Moser, E. The preparation and readiness for voluntary movement: a high-field event-related fMRI study of the Bereitschafts-BOLD response. NeuroImage 20, 404–412 (2003). 10.1016/S1053-8119(03)00291-X

61 Fang, Z. et al. Human high-order thalamic nuclei gate conscious perception through the thalamofrontal loop. Science 388, eadr3675 (2025).

62 Bianciardi, M. Brainstem Navigator [Tool/Resource.]. 1.0 edn, (The Neuroimaging Tools and Resources Collaboratory (NITRC), 2021).

63 Benjamini, Y. & Hochberg, Y. Controlling the false discovery rate: a practical and powerful approach to multiple testing. Journal of the Royal statistical society: series B (Methodological) 57, 289–300 (1995).

64 Sugihara, G. et al. Detecting Causality in Complex Ecosystems. Science 338, 496–500 (2012). doi:10.1126/science.1227079

65 Ushio, M. et al. Fluctuating interaction network and time-varying stability of a natural fish community. Nature 554, 360–363 (2018). 10.1038/nature25504

66 Wismüller, A. et al. Nonlinear Functional Connectivity Network Recovery in the Human Brain with Mutual Connectivity Analysis (MCA): Convergent Cross-Mapping and Non-Metric Clustering. Proc SPIE Int Soc Opt Eng 9417 (2015). 10.1117/12.2082124

67 Gold, J. I. & Shadlen, M. N. The neural basis of decision making. Annu Rev Neurosci 30, 535–574 (2007). 10.1146/annurev.neuro.29.051605.113038

68 Koevoet, D. et al. Effort drives saccade selection. eLife 13, RP97760 (2025). 10.7554/eLife.97760

69 Kaneko, K. Life: an introduction to complex systems biology. (Springer, 2006).

70 Conrad, M. Adaptability: The significance of variability from molecule to ecosystem. (Springer Science & Business Media, 2012).

71 Beaty, R. E., Benedek, M., Barry Kaufman, S. & Silvia, P. J. Default and Executive Network Coupling Supports Creative Idea Production. Scientific Reports 5, 10964 (2015). 10.1038/srep10964

72 Hikosaka, O. & Wurtz, R. H. Modification of saccadic eye movements by GABA-related substances. II. Effects of muscimol in monkey substantia nigra pars reticulata. J Neurophysiol 53, 292–308 (1985). 10.1152/jn.1985.53.1.292

73 Chevalier, G. & Deniau, J. M. Disinhibition as a basic process in the expression of striatal functions. Trends in Neurosciences 13, 277–280 (1990). 10.1016/0166-2236(90)90109-N

74 Suzuki, M., Pennartz, C. M. A. & Aru, J. How deep is the brain? The shallow brain hypothesis. Nature Reviews Neuroscience 24, 778–791 (2023). 10.1038/s41583-023-00756-z

75 Peters, A. J., Fabre, J. M. J., Steinmetz, N. A., Harris, K. D. & Carandini, M. Striatal activity topographically reflects cortical activity. Nature 591, 420–425 (2021). 10.1038/s41586-020-03166-8

76 Friston, K. J. et al. Event-Related fMRI: Characterizing Differential Responses. NeuroImage 7, 30–40 (1998). 10.1006/nimg.1997.0306

77 Morito, Y. & Murata, T. Accumulation System: Distributed Neural Substrates of Perceptual Decision Making Revealed by fMRI Deconvolution. The Journal of Neuroscience 42, 4891–4912 (2022). 10.1523/jneurosci.1062-21.2022

78 Holmes, A. P. & Friston, K. J. Generalisability, Random Effects & Population Inference. NeuroImage 7 (1998).

79 Mumford, J. A. & Nichols, T. Simple group fMRI modeling and inference. NeuroImage 47, 1469–1475 (2009). 10.1016/j.neuroimage.2009.05.034

80 Pedregosa, F. et al. Scikit-learn: Machine learning in Python. the Journal of machine Learning research 12, 2825–2830 (2011).

81 Virtanen, P. et al. SciPy 1.0: fundamental algorithms for scientific computing in Python. Nature Methods 17, 261–272 (2020). 10.1038/s41592-019-0686-2

82 Park, J. pyEDM [Computer software]: Python/Pandas toolset for Empirical Dynamic Modeling., <https://github.com/SugiharaLab/pyEDM> (2025).

